# Super-resolving light fields in microscopy: depth from disparity

**DOI:** 10.1101/527432

**Authors:** Ruth R. Sims, Sohaib Abdul Rehman, Martin O. Lenz, Leila Mureşan, Kevin O’Holleran

## Abstract

Single molecule localisation microscopy (SMLM) has opened a new window for imaging fluorescently labelled biological specimens. Common 3D SMLM techniques enable data collection across an axial range of 1 − 5*μ*m with high precision. Despite the success of 3D single molecule imaging there is a real need to image larger volumes. Here we demonstrate, through simulation and experiment, the potential of Single Molecule Light Field Microscopy (SMLFM) for extended depth-of-field super-resolution imaging, extracting 3*D* point source position by measuring the disparity between localizations of a point emitter in multiple perspective views.

## I. INTRODUCTION

Light field microscopy (LFM) offers single-shot, three-dimensional imaging by simultaneously collecting light from a large, continuous depth-of-field and discriminating emitter location through wavefront sampling. This incredible capability is generally achieved at a cost - sampling the wavefront and recording directional information reduces spatial resolution [1]. For instance, linear integral reconstruction of LFM data typically causes an order of magnitude loss in lateral spatial resolution when compared to a scanned system with similar optics [1]. Recent developments have utilized knowledge of the light field point spread function to recover some high resolution information, reconstructing volumes using computationally intensive deconvolution methods [2, 3]. The fast volumetric imaging capabilities of LFM have been exploited to great effect in whole-brain imaging [4, 5], but it’s intermediate spatial resolution has restricted application of the technique beyond this domain. Here we demonstrate that the fundamental limits to light field resolution can be circumvented by localizing point sources in perspective views of the specimen.

LFM is commonly implemented by placing a microlens array (MLA) in the image plane of a widefield microscope. As illustrated in Fig. 1, the MLA acts to spatially sample the wavefront of fluorescence in the image plane at a frequency determined by the microlens pitch. Each, locally filtered, wavefront is then focused onto a sub-region of a camera chip, creating a partitioned 2D space of virtual-pixels. This array of virtual pixels can then be mapped to the 4D light field, *L*(**R**, **K**), a pairing of discrete spatial (microlens) index, **R**, and discrete angular coordinate, **K**, locally within each virtual-pixel. The 3D location of a point emitter is encoded in the light field through the nature of the MLA’s effect on the incident wavefront curvature (defocus) and tilt (lateral position).

**FIG. 1.**
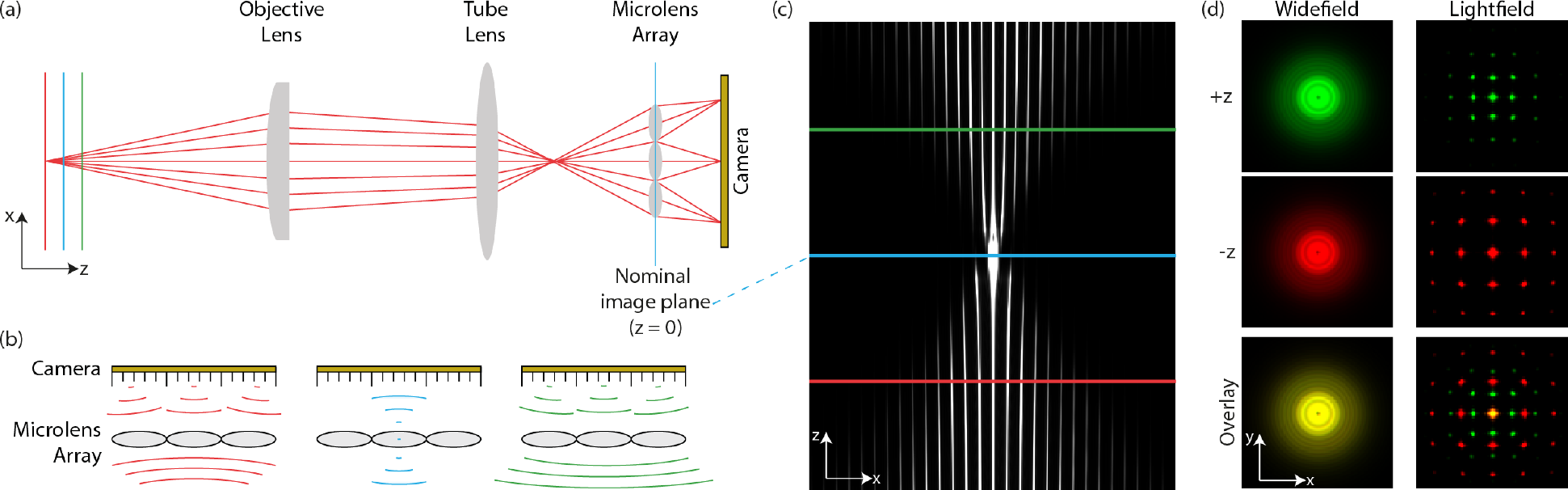
Wavefronts recorded in light field microscopes encode the three-dimensional location of point emitters. (a) Schematic diagram of the optical setup used to capture light field measurements. (b) In LFM a microlens array located at the image plane spatially samples incident wavefronts. (c) An XZ slice through the light field point spread function. (d) Comparison of different XY slices through simulated widefield and light field point spread functions. The transformation of wavefront curvature into intensity breaks the axial degeneracy of the widefield point spread function.

Wavefront curvature due to a defocused point source is symmetric around its lateral location and exhibits a sign inversion at the image plane. A result of this inversion is that the light field due to a point source maps to a unique 3*D* spatial location. Techniques exploiting the asymmetry of wavefront curvature about the object plane in order to localize point sources in 3*D* have previously been demonstrated [6], with the largest depth of field obtained using diffractive optical elements to reshape the PSF [7–10]. The implementation of LFM described here employs an off-the-shelf, refractive optical element (MLA), which offers many advantages: large spectral bandwidth for multi-colour imaging, low component cost and high photon throughput.

We have developed an algorithm capable of estimating the three-dimensional position of a point source from a light field with sub-wavelength precision. Our approach is based on an intuitive manifestation of light field shearing with defocus: parallax [11–13]. The characteristics of the technique were explored using a combination of simulations and experiments performed on a modified, inverted high NA microscope. The viability of this method was verified by observing fluorescent microbeads freely diffusing in solution and through detection of single molecule photobleaching events in immunolabeled cells.

## II LOCALIZING POINTS IN LIGHT FIELDS

To localize point emitters in light fields using parallax it is first necessary to render the light field into *perspective views*, which are generated by grouping together elements in the light field with common **K**. This is equivalent to taking a cross-section of *L*(**R**, **K**) at **K**= **K**\*. Each perspective view, *P*_**K**\*_ (**R**), is a spatial image, coarsely sampled by the microlens pitch and (through selection of **K**\*) filtered by a particular wavefront tilt (or, equivalently, a ray direction). The image of the point source in each *P*_**K**\*_ is localized, by fitting a two-dimensional Gaussian function, to yield a measurement of source location, represented as a delta function, *δ*(**r**−Δ**r**, **k**− **K**\*), in the (now continuous) light field 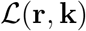. Localization is performed across all detected emitters in all perspective views resulting in a set of detected localizations:

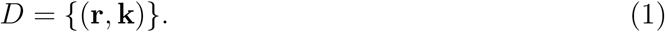

This set, comprising of all detected localizations, is then mapped into a collection of subsets, *S*, which is an exact cover of *D*, through nearest neighbour tracking in adjacent perspective views. Each subset in *S* represents a single detected emitter, identified by grouping multiple localizations of the same source from different perspective views. The sign function of each point emitter (that is, whether the emitter is above or below the object plane) is given by the direction of the tracks along the principle axes: sgn(Δ*r*). In the absence of other aberrations, the circular symmetry of defocus constrains these grouped localizations to the surface of a cone in (**r**, *k*_*r*_) where *k*_*r*_ = |**k**|. This cone, a mapping of parallax across all perspective views, contains all relevant 3D positional information: its gradient, *α*(|*z*|) and apex **r**_0_ respectively encoding the lateral location and the modulus of the axial location of the point-source:

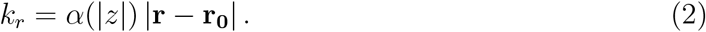

The global axial position of each point source is then retrieved by multiplication with sgn(Δ*r*). This procedure is summarized in Algorithm 1 with key steps being illustrated in Fig. 2.

### Algorithm 1 Process for 3D localization of point emitters in captured, rectified light fields.

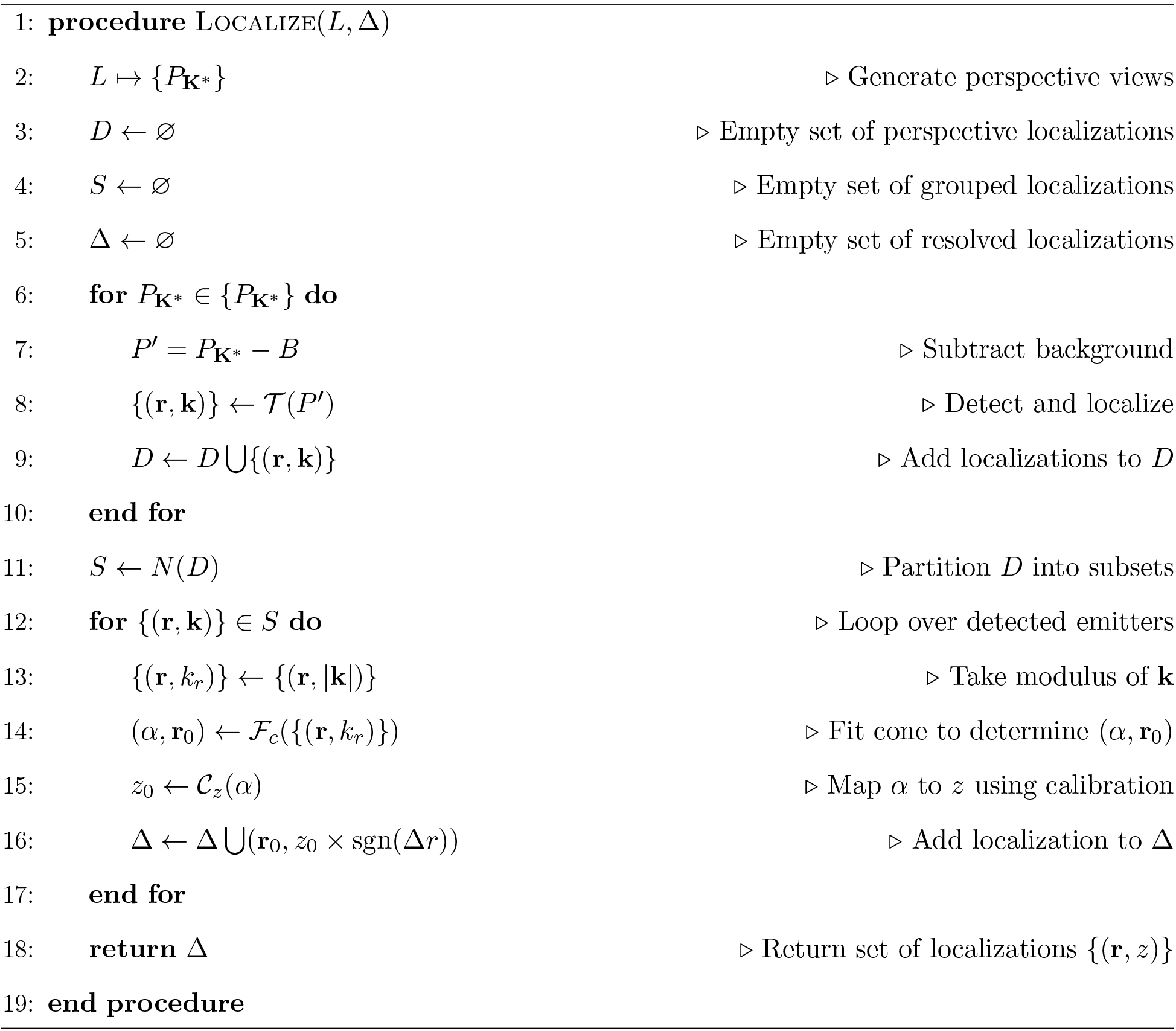

**FIG. 2.**
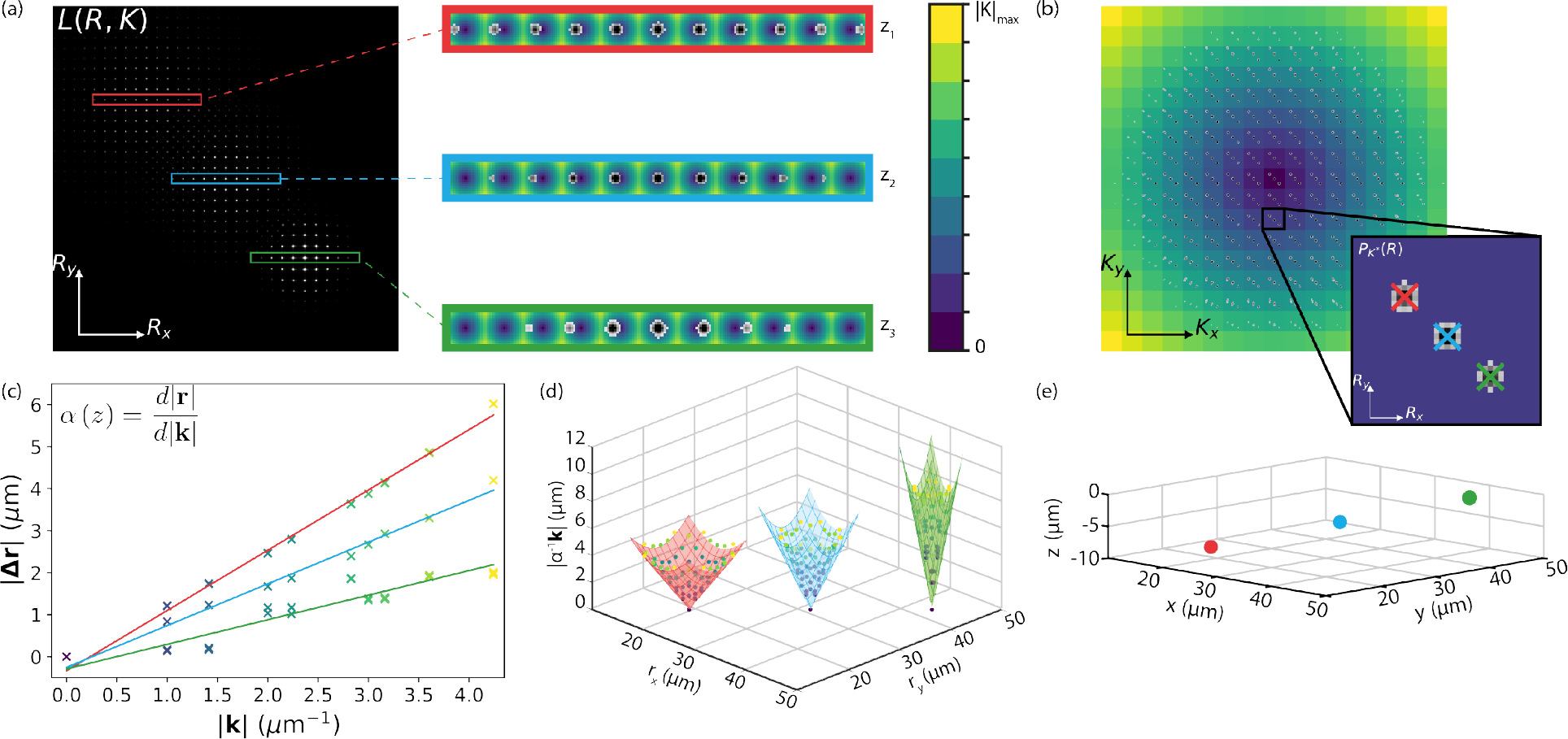
Recovery of super-localized three-dimensional point source locations from a light field measurement. (a) Simulated light field measurement due to three point sources at distinct locations in object space. (b) Point sources are localized in each perspective view. (c) The position of the point source in each perspective view is linearly proportional to its axial position. (d) The localizations corresponding to each emitter are constrained to the surface of a cone in (**r**, *k*_*r*_). The 3*D* position of the emitter is encoded in the gradient, *α*(*z*) and apex **r**_0_ of this cone. For more details refer to part 2 of the Supplementary Information. (e) *z* is estimated from *α* (*z*) using an experimentally recorded calibration curve.

It is well established that aliased, high-frequency information can be used to generate super-resolved images (with respect to the lenslet sampling rate), indeed this is responsible for the improved resolution of deconvolved light field images [2, 3, 12]. The method presented here exploits the fact that in the case of point sources, these sub-pixel shifts can be measured directly. In effect, this approach reduces the cumbersome 5D light field point spread function to just two parameters which define the form of a right-circular cone: *α* and **r**_0_. This is a direct result of the phase-space measurement of a point source being constrained to a hyperplane in 4D, fully characterized by its intercept and gradient [14]. Whilst the redundancy of a light field measurement due to a point source is recognized [15], this geometric approach combines information from these redundant measurements in order to determine the three-dimensional position of a point source with unprecedented precision as compared with existing light field methods. Instead of trading spatial resolution for angular information, this approach is capable of three-dimensional point source localization with a precision far smaller than the diffraction limited spot of the corresponding widefield microscope.

## III. SIMULATING LIGHT FIELDS

The geometric approach to three-dimensional localization was tested by estimating the position of point emitters in simulated light field datasets. The optical model used to perform simulations was designed to closely replicate the corresponding experimental system. The simulations were based on some key assumptions: monochromatic, isotropic fluorescent emission from point sources with no scattering and scalar diffraction. The full microlens array was generated by convolving its transmittance function with a two dimensional Dirac comb. The phase profile of the microlens array was multiplied with the widefield image of the point source and propagated to the camera (refer to Section 2 of the Supplementary Information for full details).

Using this model, light field measurements corresponding to a point source axially displaced over a 30 *μ*m range with 100 nm step size, were generated using parameters matched to those of the experimental setup. Results plotted in Fig. 3 demonstrate that the gradient of disparity, *α*, exhibits non-linear behaviour with axial point source position. This is a consequence of local phase curvature within a microlens. Since localization indirectly identifies the most significant tilt component of the local wavefront, any deviation from piece-wise linearity results in a non-linear relationship between *α* and *z*. As the wavefront expands with defocus, piece-wise decomposition across microlenses becomes a closer approximation to the true wavefront and *α*(*z*) tends to a linear relationship. The most extreme deviation from linearity occurs within 1*μ*m of the image plane. The precision of position estimation is poor in this region since both localization in **P_k_** and solving Equation 2 become difficult. *α*(*z*) remains monotonic in-spite of these non-linearities and, most importantly, can be seen to act as an excellent axial location discriminator.

**FIG. 3.**
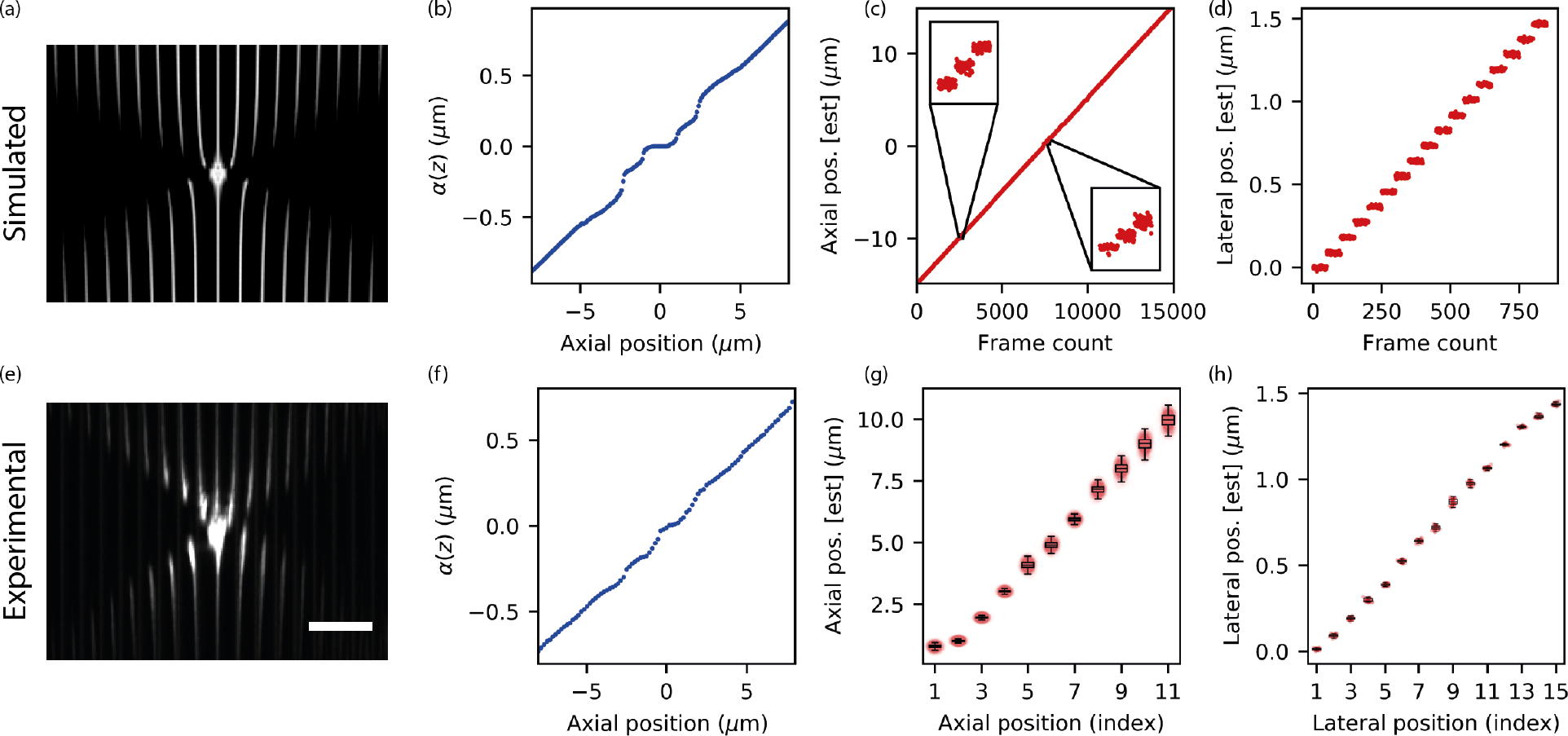
Sub-diffraction limited localization precision demonstrated with simulated and experimental light field data. (a-d) Results of simulations based on the optical model described in Section III. (e-h) Experimental results for characterizing our light field microscope using 100 nm fluorescent beads as a test sample. (a) XZ slice of the light field point spread function (max projection). (b) Calibration curve generated by calculating *α* (*z*) for simulated light fields corresponding to point sources at different axial positions along the optical axis between *±*8 *μ*m. (c) Calculated axial position of a simulated emitter scanned, along the optical axis, between *±*15 *μ*m with a step size of 100 *nm*. (d) Calculated lateral position of an emitter scanned over 1.5 *μ*m (pitch of the lenslet array used in the experiments) with a step size of 100 nm. 50 images were generated at each position in (c) and (d). (e) XZ slice of the experimental light field point spread function. (f) Calibration curve generated by scanning beads between *±*8 *μ*m with a step size of 100 nm. 10 images were captured at each axial position and the calibration curve was generated from the mean axial position of a bead localized using Algorithm 1. (g) Calculated axial position of a bead scanned over a distance of 12 *μ*m (on one side of the focal plane) with a step size of 1 *μ*m. Box plot was generated from mean and standard deviation of the axial position calculated from 500 images of the bead at each position. (h) Calculated lateral position of a bead scanned over a distance of 1.5 *μ*m with a step size of 100 nm. Box plot was generated from mean and standard deviation of the lateral position calculated from 30 images captured at each position. (a) and (e) are displayed with the same pixel size. Scale bar in (e) corresponds to 5 *μ*m.

**FIG. 4.**
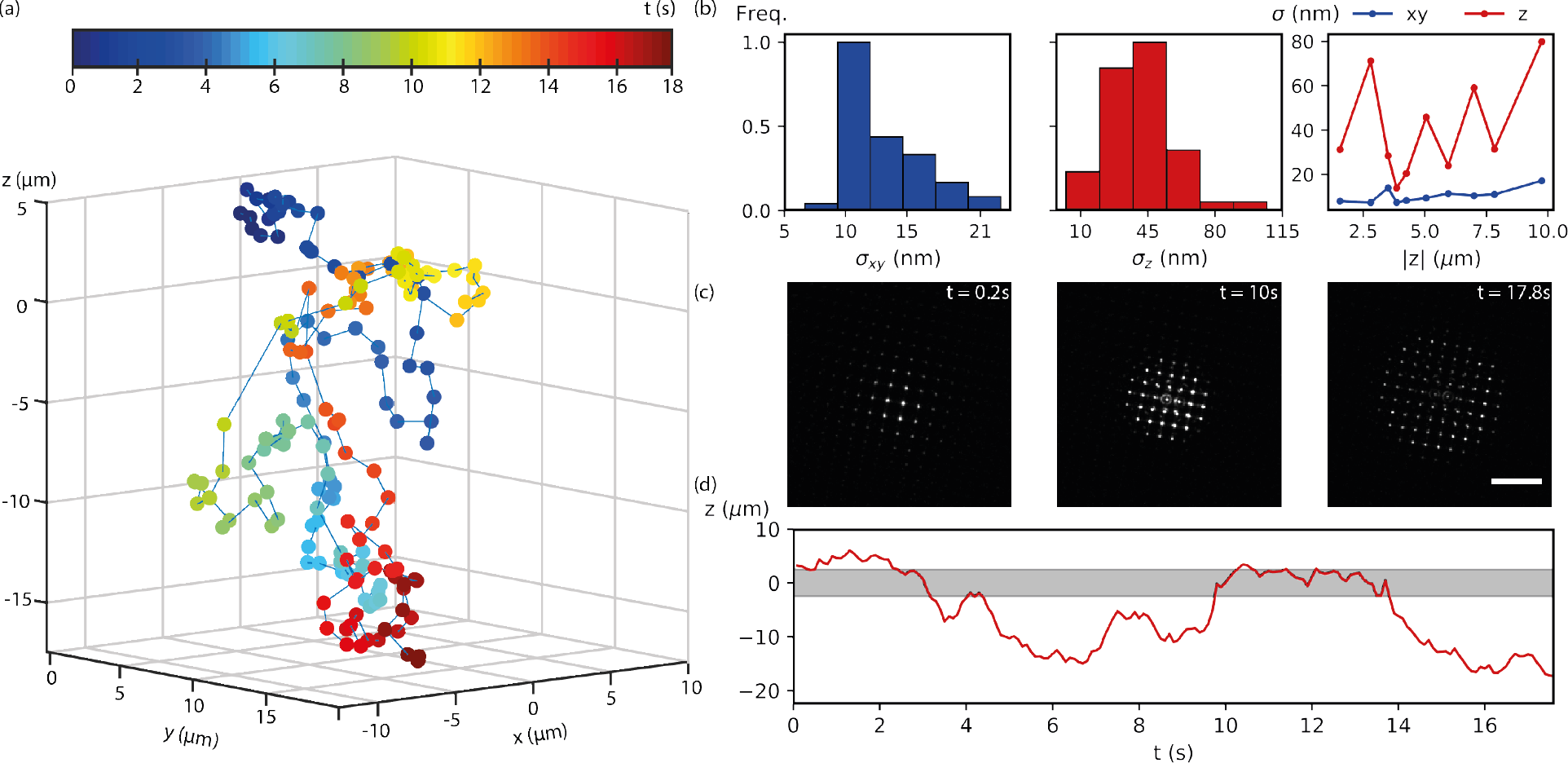
Tracking a freely diffusing 100 nm fluorescent bead over an extended depth of field with light field microscopy. (a) Estimated 3*D* trajectory of a representative 100 nm fluorescent bead undergoing Brownian motion in water. Each point is colour coded as a function of times. The trajectory was calculated by estimating the 3*D* location of the bead in each frame using the procedure summarized in Algorithm 0. (b) Histograms of lateral and axial precision throughout the depth of field. (c) Raw frames of bead at different time points. Scale bar represents 10 *μ*m. (d) Axial location of bead as a function of time. Grey region indicates the depth of field achieved by common single molecule localization microscopy techniques. Precision estimates for the diffusing bead were estimated by comparison with calibration data with comparable numbers of photons. Further details may be found in Section X of the Supplementary Information.

## IV. EXPERIMENTAL RESULTS

The concept of localizing point sources in light fields was experimentally validated by modifying an inverted widefield microscope to incorporate light field detection. Proof-of-concept experiments were performed by positioning an approximately *f*-number matched MLA in the image plane. *f*-number matching the MLA and the numerical aperture of the objective lens eliminates cross talk between virtual-pixels. A calibration curve was generated by axially scanning a 2*D* sample of 100 nm diameter fluorescent beads over an axial range of 16 *μ*m, with step size of 100 nm and calculating *α*(*z*) at each position. The results plotted in Fig. 3(f) closely match the corresponding curve generated based on simulated light fields of Fig. 3(b). The localization precision of LFM was measured by capturing 500 images at each axial position between 0 and 12 *μ*m with 1 *μ*m separation. The precision was calculated as the standard deviation of all localizations at each position. Since, theoretically, *α* (*z*) is identical either side of the object plane, it was only necessary to capture data from one side of the object plane. This allowed the precision to be calculated from a large number of measurements without significant photobleaching between initial and final positions.

The capabilities of the system with respect to data acquisition across an extended depth of field of 25 *μ*m, were demonstrated by imaging 100nm fluorescent beads freely diffusing in water. Data from a typical bead, tracked for 18 s, at 100 ms intervals is summarized in Fig. 3. The bead was localized in each frame using the workflow summarized in Figure 2. *α* (*z*) was converted into *z* position using the experimental calibration curve plotted in (f) of Fig. 3.

Two experiments were performed to demonstrate the feasibility of single molecule imaging using light field microscopy. Membrane (TCR) proteins labelled with Cage-552 dye were imaged in fixed T-cells, using 561 nm illumination and 20 ms exposure time. Histones labelled with Alexa-647 were imaged in drosophila spermatocytes, using 638 nm illumination and 50 ms exposure time. Full details of the sample preparation protocols for both sets of experiments are presented in the Supplementary Information. Typical fluorescent traces from each set of experiments are plotted in Fig. 5. These exhibit discrete signal levels, characteristic of single molecule photobleaching events [16].

**FIG. 5.**
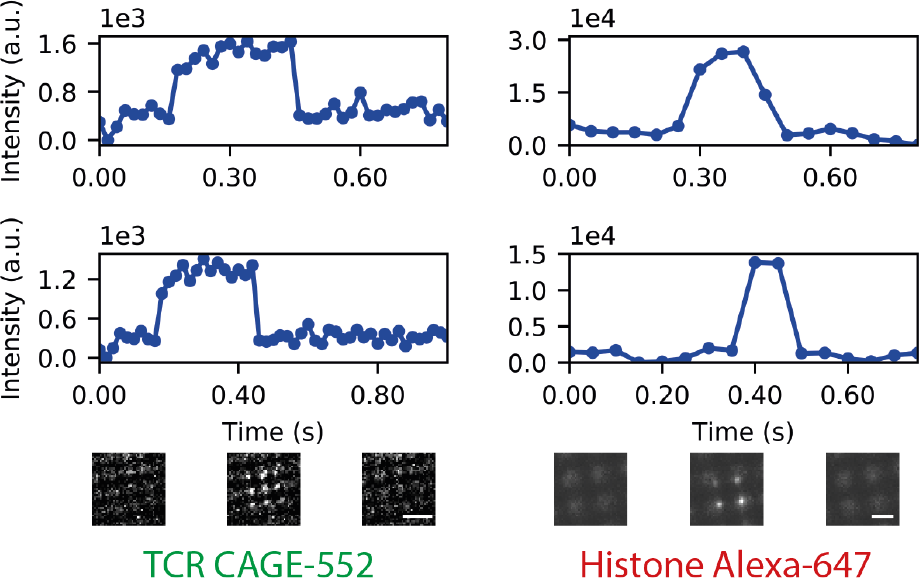
Single molecule photobleaching events captured using light field microscopy. Photobleaching curves of CAGE-552 fluorophores imaged on the surface of fixed T cells and histones from drosophila spermatocytes labelled with Alexa-647 imaged with light field microscopy. In each case integrated intensity is plotted as a function of time. Camera frames with background subtracted are also shown at different time points. Scale bars represent 1.5 *μ*m.

## V. DISCUSSION

These results indicate that light field microscopy is a technique with significant potential for imaging single molecules throughout an extended depth-of-field. Simulations and experimental data demonstrate that sub-diffraction limited localization precision over a continuous 10^4^ *μ*m^3^ volume is readily achievable. While proof-of-principle experiments were carried out using fluorescent beads which emit relatively large numbers of photons, the viability of light field microscopy as a single molecule imaging technique was demonstrated by imaging single labelled proteins in fixed T-cells and histones in fixed drosophila spermatocytes. In these experiments, single molecule bleaching events were observed throughout a 10 *μ*m axial volume.

These results were acquired on a prototype system incorporating a sub-optimal, off-the-shelf, microlens array. It is anticipated that modification of the MLA characteristics along with further development of the algorithm will facilitate the application of light field microscopy to imaging single molecules throughout the entire volume of a mammalian cell. In order to reach this goal, the localization precision in low photon number and scattering regimes must be improved.

Such improvements could be readily achieved by a simple modification of the experimental setup, namely optimizing the microlens pitch according to the required axial range. Using smaller lenses would result in better lateral precision, particularly at the focal plane, but suffer in terms of shot noise due to the distribution of photons across an increased number of pixels. Another simple modification would be to position the microlens array in a plane conjugate to the back focal plane of the microscope objective. In this configuration, the camera would directly capture perspective views [17, 18], the number of which would be dictated by the number of microlenses spanning the diameter of the pupil and the localization precision would be dictated by the effective camera pixel size as is the case in typical 2*D* single molecule imaging experiments [19].

Furthermore, algorithms capable of exploiting the over-determined nature of each light field measurement ought to be investigated. The redundancy of this measurement arises since the light field due to a single point source is comprised of > *n*^2^ data points, where *n* refers to the number of camera pixels spanned by each microlens. In typical configurations of light field microscopes, *n* ranges between 15 and 20 pixels [20] and, correspondingly, the inverse problem of estimating the three dimensional location of a point source is extremely over-determined. On the other hand, since typical imaging volumes contain significantly more voxels than the number of measured pixels [21], reconstruction of the entire volume by Richardson-Lucy deconvolution is under-constrained. As a result, although deconvolution approaches have been successful, the resolution that can be obtained is fundamentally limited [22]. Whilst the geometric approach presented here is more tractable than deconvolution, sparsity of emitters in the spatial domain is necessitated by the use of localization algorithms. Emitter sparsity is an inherent feature of single molecule imaging experiments. Furthermore, this approach allows for simple system calibration, where the effect of aberrations are absorbed into the light field, manifested as a change in the rate of light field shearing as a function of defocus. On the other hand, to generate high-fidelity images using deconvolution, any deviations from the ideal optical model must be identified and specifically accounted for.

It is anticipated that this technique will find application for imaging in ballistic scattering regimes such as biological tissue since previous studies have exploited the redundancy of the light field measurement to mitigate the effects of volumetric scattering [15, 23]. Identifying the appropriate balance between measurement redundancy and signal-to-noise ratio will be the subject of future work.

## VI. CONCLUSION

The viability of SMLFM for sub-diffraction localization has been demonstrated through extraction of depth information through disparity of localizations of point sources in perspective views. Simply using stock microlens arrays without further optimization allowed 50nm precision. Taking this concept to genuine whole-cell imaging will require improvement of the localization pipeline to exploit the over-determined nature of the light field PSF. Exploiting this property to improve precision at low photon numbers would position SMLFM as one of the most attractive developments in high-resolution microscopy.

## AUTHOR CONTRIBUTIONS

RRS and SA designed and performed simulations and experiments. RRS designed and implemented the lightfield rectification algorithm. LAM lead the data analysis in collaboration with RRS and SA. Experimental apparatus was implemented by MOL and SA. KOH and RRS, wrote the manuscript. KOH conceived and managed the project. All authors contributed to figure design and edited the final manuscript.

## ACKNOWLEDGMENTS

The authors thank Dr Stephen F. Lee for useful discussions and Dr Rob White and Dr Alexander R. Carr for providing labelled drosophila spermatocyte cells and T-cells respectively.

## SUPPLEMENTAL DOCUMENTS

See Supplement 1 for supporting content.

## REFERENCES

[1] M. Levoy, R. Ng, A. Adams, M. Footer, and M. Horowitz, ACM Transactions on Graphics (TOG) 25, 924 (2006).

[2] M. Broxton, L. Grosenick, S. Yang, N. Cohen, A. Andalman, K. Deisseroth, and M. Levoy, Optics Express 21, 25418 (2013).

[3] T. E. Bishop, and P. Favaro, IEEE Transactions on Pattern Analysis and Machine Intelligence 34, 972 (2012).

[4] R. Prevedel, Y.-G. Yoon, M. Hoffmann, N. Pak, G. Wetzstein, S. Kato, T. Schrödel, R. Raskar, M. Zimmer, E. S. Boyden, et al., Nature Methods 11, 727 (2014).

[5] O. Skocek, T. Nöbauer, L. Weilguny, F. M. Traub, C. N. Xia, M. I. Molodtsov, A. Grama, M. Yamagata, D. Aharoni, D. D. Cox, et al., Nature Methods, 1 (2018).

[6] B. Huang, W. Wang, M. Bates, and X. Zhuang, Science 319, 810 (2008).

[7] S. R. P. Pavani, M. A. Thompson, J. S. Biteen, S. J. Lord, N. Liu, R. J. Twieg, R. Piestun, and W. Moerner, Proceedings of the National Academy of Sciences 106, 2995 (2009).

[8] P. Bon, J. Linarès-Loyez, M. Feyeux, K. Alessandri, B. Lounis, P. Nassoy, and L. Cognet, Nature Methods (2018).

[9] Y. Shechtman, L. E. Weiss, A. S. Backer, S. J. Sahl, and W. E. Moerner, Nano Letters 15, 4194 (2015), pMID: 25939423, https://doi.org/10.1021/acs.nanolett.5b01396.

[10] A. von Diezmann, Y. Shechtman, and W. Moerner, Chemical Reviews 117, 7244 (2017).

[11] A. Isaksen, L. McMillan, and S. J. Gortler, Proceedings of the 27th annual conference on Computer graphics and interactive techniques, 297 (2000).

[12] E. H. Adelson, and J. Y. A. Wang, IEEE Transactions on Pattern Analysis & Machine Intelligence, 99 (1992).

[13] R. Ng, M. Levoy, M. Brédif, G. Duval, M. Horowitz, P. Hanrahan, et al., Computer Science Technical Report CSTR 2, 1 (2005).

[14] D. Dansereau and L. Bruton, Proceedings of the 2004 International Symposium on Circuits and Systems, 2004 3, III (2004).

[15] H.-Y. Liu, E. Jonas, L. Tian, J. Zhong, B. Recht, and L. Waller, Optics express 23, 14461 (2015).

[16] C. Liesche, K. S. Grußmayer, M. Ludwig, S. Wörz, K. Rohr, D.-P. Herten, J. Beaudouin, and R. Eils, Biophysical journal 109, 2352 (2015).

[17] L. Cong, Z. Wang, Y. Chai, W. Hang, C. Shang, W. Yang, L. Bai, J. Du, K. Wang, and Q. Wen, Elife 6, e28158 (2017).

[18] G. Scrofani, J. Sola-Pikabea, A. Llavador, E. Sanchez-Ortiga, J. Barreiro, G. Saavedra, J. Garcia-Sucerquia, and M. Martínez-Corral, Biomedical optics express 9, 335 (2018).

[19] R. E. Thompson, D. R. Larson, and W. W. Webb, Biophysical journal 82, 2775 (2002).

[20] M. Shaw, M. Elmi, V. Pawar, and M. A. Srinivasan, Biomedical optics express 7, 2877 (2016).

[21] T. Nöbauer, O. Skocek, A. J. Pernía-Andrade, L. Weilguny, F. M. Traub, M. I. Molodtsov, and A. Vaziri, Nature methods 14, 811 (2017).

[22] I. Kauvar, J. Chang, and G. Wetzstein, IEEE International Conference on Computational Photography, 1 (2017).

[23] N. C. Péegard, H.-Y. Liu, N. Antipa, M. Gerlock, H. Adesnik, and L. Waller, Optica 3, 517 (2016).

