## Supplementary text and figures for "Super-resolving light fields in microscopy: depth from disparity"

### Super-resolving lightfields in microscopy: depth from disparity
### Supplementary Information

Ruth R. Sims,[1] Sohaib Abdul Rehman,[1] Martin O. Lenz,[1] Leila Mureşan,[1] and Kevin O'Holleran[1]
[1] *Cambridge Advanced Imaging Centre, University of Cambridge, Downing Site, Cambridge, CB2 3DY*

#### CONTENTS

I. Methods and Experimental Verification 1
A. Experimental Apparatus 1
B. Alignment and Rectification 1
C. Localization Precision 2
D. Background Estimation 2
E. Detection / Localization 3
F. Localizing Multiple Emitters 3

II. Theoretical Background 4
A. The light field due to a point source 4
B. Optical Imaging Model 5

III. Sample preparation 6
A. T Cell Labelling Protocol 6
B. Spermatocytes Labelling Protocol 6

References 6

#### I. METHODS AND EXPERIMENTAL VERIFICATION

##### A. Experimental Apparatus

All experimental data presented in this work was acquired on a customized, inverted microscope (Leica DM IRE2), shown in Figure 1. Illumination for excitation and activation was provided by a set of lasers (405 nm, 488 nm, 561 nm and 638 nm) housed in an Omicron LightHUB. The output from each laser was coupled into a single mode optical fibre before being collimated into an 8 mm diameter beam, using an off-axis parabolic mirror (Thorlabs RC08APC-P01) in order to minimise chromatic aberrations. The two mirrors labelled $M1$ and $M2$ in Figure 1 were adjusted to dictate the necessary beam deflection for sample illumination by either 'epi' or HILO. This beam was focused onto the back focal plane of a $60x$, 1.3NA Silicone oil immersion objective (Olympus, UPLSAPO60XS) using a 250 mm focal length lens, $L1$, resulting in a 96 $\mu$m diameter beam (FWHM) at the sample. A quad-band filter (Chroma ZT405/488/561/640rpc) was placed between $L1$ and the objective to separate excitation and activation beams from the fluorescence. Samples were placed on a 3 axis piezo-driven stage (Nanos, LPS 30-60-1-VX-S-N-XY and LPS 30-30-1-VX-S-N-XYZ) to allow 3-dimensional movement with a resolution down to 10 nm.

Fluorescence was collected using the same objective and focused using a 200 mm focal length lens, $L2$. The f-number matched microlens array (RPC Photonics MLA S100-f21, pitch = 100$\mu$m, $f$ = 2.1mm) was placed in the image plane of the microscope. The back focal plane of the microlens array was imaged on an sCMOS camera (Hamamatsu Flash 4.2) using a 1 : 1 relay system (Lenses L3 and L4, $f$ = 125 mm focal length).

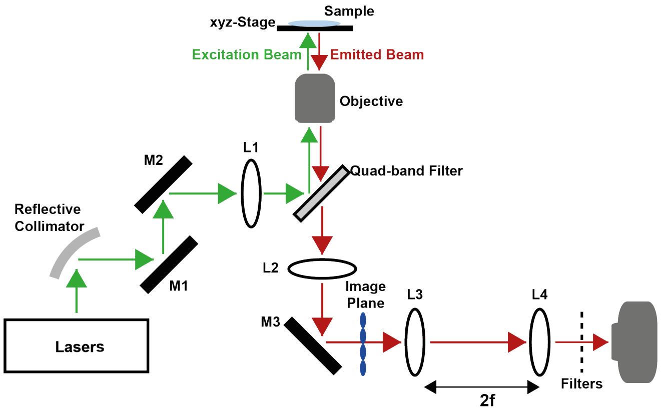


FIG. 1. Schematic diagram of experimental setup. M - Mirror, L - Lens, 2f - distance between lenses is sum of their focal lengths.

##### B. Alignment and Rectification

Similarly to `http://graphics.stanford.edu/software/LFDisplay/lfmintro//`, the microlens array was placed in the image plane of a widefield microscope using the following procedure. Firstly, collimated light was directed through the microscope objective. The microlens array was translated about the approximate location of the image plane until the square profile of the microlenses could be seen on the camera. Next, the camera was axially shifted to image the back focal plane of the microlens array. At this position a set of focused spots was observed on the detector due to the microlenses focusing the collimated beam. The axial location of the microlens array was verified by imaging a thin fluorescent layer. For a well aligned system, the corresponding image is composed of uniformly intense circles since in this configuration each microlens is imaging the back focal plane of the microscope objective.

During each experiment a calibration image was acquired. Images of a thin fluorescent layer or collimated light directed through the microscope objective were used. These images were used to extract rectification parameters (microlens pitch and rotation and translation of microlens array with respect to the camera). Briefly, the rotation was identified by measuring the orientation which maximized the variance of the radon transform of the calibration image. The pitch was calculated by localizing harmonics in Fourier space with sub-pixel resolution. Finally the translation was calculated by generating a grid of points spanning the calibration image using the estimated rotation and microlens pitch. This grid is translated a distance of 1.5 microlenses. At each position a metric is calculated (for instance, for f-number matched microlens arrays a suitable metric is the sum of intensity at each grid point). The centroid of this metric corresponds to the translation of the microlens array.

These parameters were subsequently used to rectify all data acquired experimentally before generating the perspective views.

##### C. Localization Precision

The axial and lateral precision of the method was determined under different experimental conditions using 100 nm diameter fluorescent beads scanned over an axial range of 12 $\mu$m, with a step size of 1 $\mu$m. 500 images were captured at each position and the precision was calculated from the standard deviation of localized beads' positions over these frames. Imaging was carried out at different laser powers, which resulted in different number of photons captured at the camera i.e. $16 \times 10^3$, $39 \times 10^3$, $47 \times 10^3$, $16 \times 10^3$, and $2.93 \times 10^5$. Figure 2 shows the results of the analysis.

(A) of Figure 2 shows the position of a fluorescent bead as a function of image number (a proxy for stage position). This graph shows the algorithm is capable of calculating the axial position, correctly separated by 1$\mu$m for most of the axial range. As anticipated, the model breaks down within 1 $\mu$m of the focal plane of the objective. The precision decreases with increasing axial displacement from the focal plane due to the spread of signal over a larger number of microlenses. (B) and (C) demonstrate that decreasing numbers of photons leads to deterioration of the localization precision. The localization precision varies from 50 nm to 200 nm, at a depth of 7 $\mu$m, for $5.6 \times 10^5$ and $30 \times 10^3$ photons respectively.

(D) of Figure 2 shows lateral and axial localisation precision as a function of different positions within a $\mu$-lens at a depth of 4.5 $\mu$m. We found the precision to be independent of the position within a $\mu$-lens as the PSF covered a similar number of $\mu$-lenses. The above analysis shows that the localisation precision varies as a function of the number of photons and the axial position of an emitter. The precision was independent of the position of the emitter within a lens. This is also evident from simulation results shown in Figure 3. The figure was generated by laterally scanning a simulated emitter over the pitch of the lenslet array, for multiple axial positions.

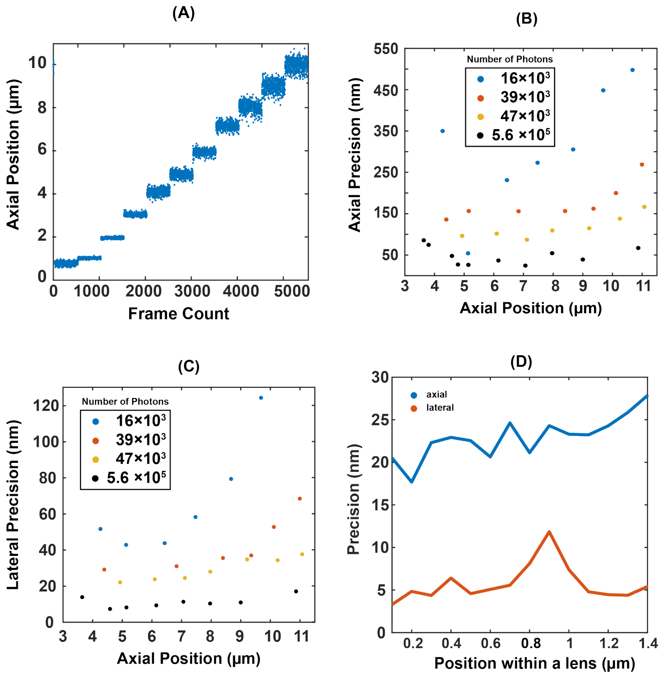


FIG. 2. (A) Caclulated axial position (using Algorithm 1) of a fluorescent bead scanned over 12 $\mu$m with a step size of 1 $\mu$m. (B-C) Axial and lateral precisions calculated from the standard deviation of the axial position calculated from multiple frames at each position of the bead. (D) Axial and lateral precisions at different positions within a micro lens

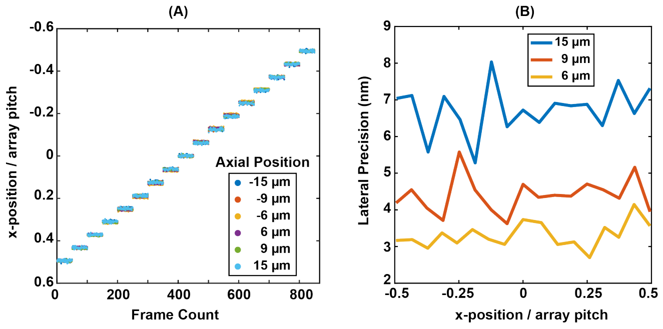


FIG. 3. Simulation results for localising an emitter scanned laterally over the lenslet array pitch for multiple axial positions

##### D. Background Estimation

The background of single shot three-dimensional imaging techniques tends to be higher than that of two dimensional counterparts. The two contributing factors to this high background are:

1. By design, the spatial frequency content of a 3D point spread function does not degrade as rapidly

2. A larger sample volume is typically illuminated.

A workflow for estimating background in light field datasets was developed, this is summarized in Figure 4.

The background was estimated in $P_k$ rather than $\mathcal{L}(\mathbf{k}, \mathbf{r})$ since the microlens array acts to focus background fluorescence into a series of sharp foci - not locally dissimilar to the light field point spread function. The fact that the light field signatures of signal and background occur at similar spatial scales makes accurate background estimation in $\mathcal{L}(\mathbf{k}, \mathbf{r})$ difficult. On the other hand, in the sub-aperture domain, the background signal appears in each view as the smoothly-varying signal familiar to many fluorescence datasets. An existing ImageJ plugin [1] was used to estimate and subsequently subtract the background contribution in each $P_k$ prior to detection and localization.

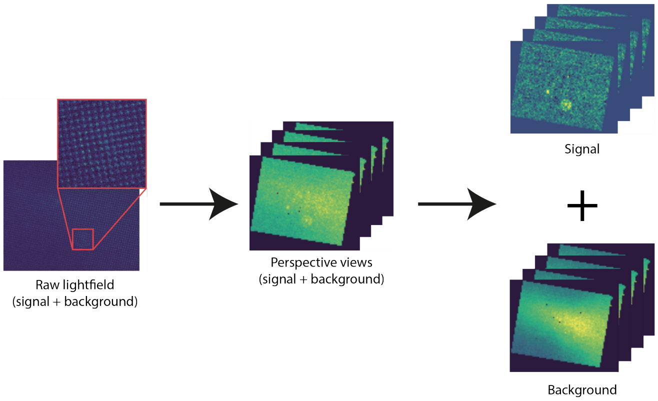


FIG. 4. Background estimation in light field datasets.

#### E. Detection / Localization

The lateral position of a point source can be determined by localization in the central perspective view ($k = 0$) since this view samples the radiance directly. However, in general, the central view has the highest background (this corresponds to the pixel the microlenses focus collimated [background] fluorescence to. In addition, this approach does not capitalize on the redundancy of a light field measurement - for the experimental configuration in Section I B this would correspond to utilization of $1/17^2$ of the total information captured.. Fitting a cone to the position of emitters in different perspective views solves these problems and gives high precision. Figure 5 shows results for calculating lateral position of the simulated light field PSF ($z = 15\,\mu$m) by direct localization in the central view and by cone fitting. As expected, localization precision was significantly better (around 3 times) in the latter case.

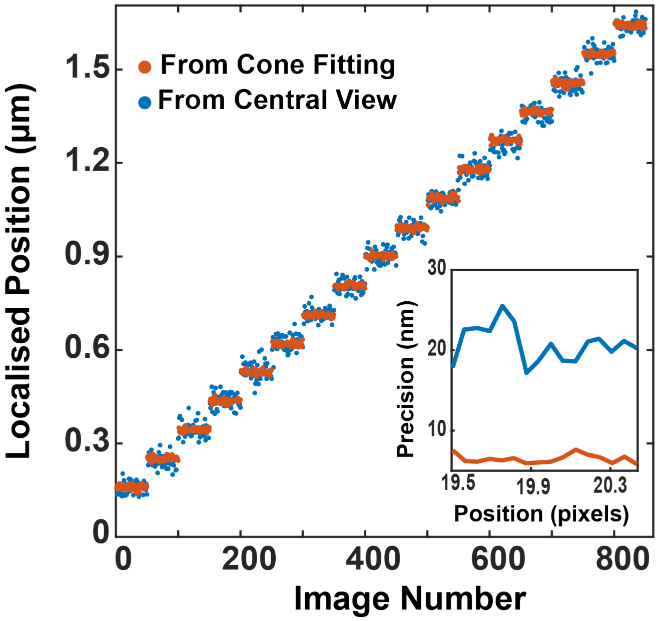


FIG. 5. Lateral position calculations, for a simulated emitter, from central view and by cone fitting.

#### F. Localizing Multiple Emitters

To calculate disparity it is important to group localisations corresponding to a specific emitter in different sub-aperture views. This was achieved by tracking localisations between perspective views. The method is demonstrate din Figure 6.

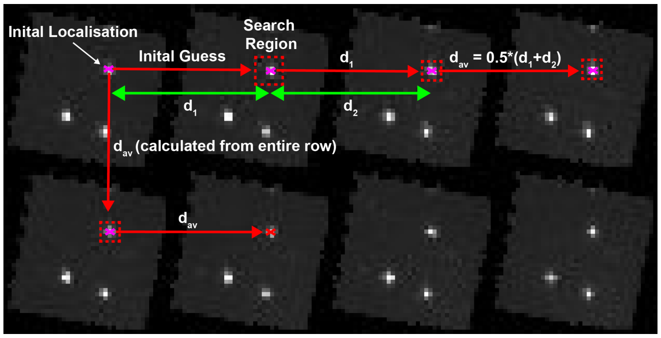


FIG. 6. Principle of our localization and tracking algorithm for multiple emitters in the field of view

The processs is carried out as follows:

1. Start by localising spots in the top left view. Let this be view 1.

2. Localise spots in the neighbouring view (along row or column), view 2, at a distance, $d_0$ (which is the initial user defined value), from localisations in view 1.

3. Calculate the distance between positions of the

emitter in views 1&2. Let this distance be $d_1$ and it depends on the axial position of the emitter.

4. Search for spots in view 3 at a distance $d_1$ from positions calculated in view 2 and perform spot localisation. Let the distance between positions of the emitter in views 2&3 be $d_2$. In the absence of aberration, we found disparity to be the same between any two views, so $d_1 \approx d_2$ (see the next section for more details).

5. In the next view, search for spots at a distance $d_{av} = 0.5 \times (d_1 + d_2)$ and repeat the process for the entire row. Although not necessary, due to constant disparity between neighbouring views, but we updated $d_{av}$ after localisation in each view.

6. Once at the end of the row, move to the next row and look for spots at a distance $d_{av}$ from localisations in view 1 (Figure 6).

7. Repeat the above mentioned steps for all the rows

Through the above steps we combined localisation and tracking to group images of the emitter in different views, when multiple emitters were present in the FOV. This method was sufficient in cases where emitters were well-separated (e.g. Figure 7). In order to localize more densely labelled samples, more sophisticated methods must be developed. This will be the subject of future work.

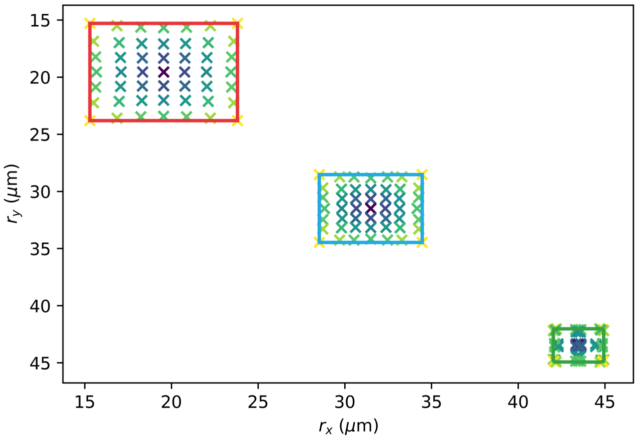


FIG. 7. The localizations of images of point emitters in each perspective view belong to distinct clusters in the case of well-separated emitters. In such regimes, nearest neighbour analysis is sufficient for clustering localizations.

#### II. THEORETICAL BACKGROUND

This section provides some mathematical intuition as to the reason why localizations due to a single point emitter are constrained to the surface cone. The optical model used to generate simulated light fields is also described in further detail than in the main text.

##### A. The light field due to a point source

All measurements in a rectified light field are constrained to satisfy the following general equation:

$$ar_x + br_y + ck_x + dk_y + C = 0 \tag{1}$$

Consider the light field due to a single isotropic fluorescent point source located at $\mathbf{P} = P\left(P_x, P_y, P_z\right)$ in object space. Only a subset of measurements defined by Equation 1 correspond to the rays due to a single point source. This subset may be identified by simple geometric considerations, assuming that the different components of the light field $\vec{k}$ and $\vec{r}$ are measured on parallel planes, between which there are no occlusions.

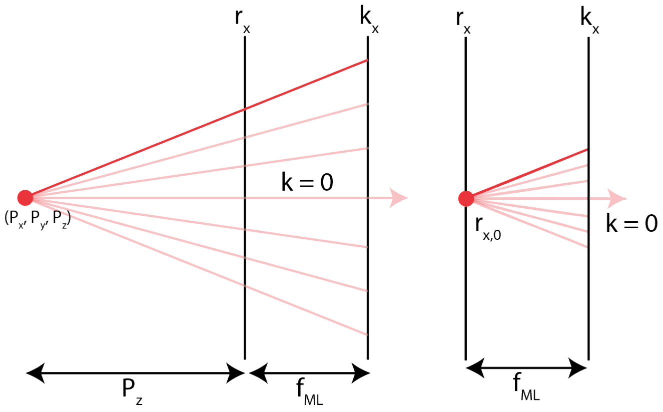


FIG. 8. The light field due to a isotropic point emitter lies on a unique plane in 4D defined entirely by its orientation and intercept.

A $2D$ slice along $k_x$ and $x$ is shown in Figure 8. It is clear that any given measurement on the $(k_x, k_y)$ plane due to the point source at $P$, can only intersect a single point on the $(x, y)$ plane. Hence, an $(x, k_x)$ slice of the light field takes the form of a line. By similar triangles, it is possible to see that the equation of this line is:

$$\left(1 + \frac{f_{ML}}{P_z}\right) r_x - k_x = \frac{P_x f_{ML}}{P_z} \tag{2}$$

Similarly for $k_y$ and $r_y$:

$$\left(1 + \frac{f_{ML}}{P_z}\right) r_y - k_y = \frac{P_y f_{ML}}{P_z} \tag{3}$$

All rays emanating from a single point source are constrained to simultaneously satisfy Equations 2 and 3. This condition means that the light field due to a point source is a plane defined by the intersection of the hyperplanes described in Equations 2 and 3. Another important characteristic of the light field due to a point source is identified by calculating the normal vectors to the hyperplanes defined by Equations 2 and 3. The normal vector is similar for the two hyperplanes, and it may be seen that $\frac{dr_x}{dk_x} = \frac{dr_y}{dk_y} = 1 + \frac{f_{ML}}{P_z}$. The orientation of the plane defined by the intersection of the hyperplanes is uniquely defined by the $z$ position of the point source $P_z$.

A plane is fully parametrized by it's orientation and intercept. According to simple physics considerations, the intercept of the plane can be seen to correspond to the transverse coordinates of the point source $(P_x, P_y)$. Figure 8 shows a point located at the object plane which is defined as the plane $P_z = 0$, such that $P_x, P_y = r_{x,0}, r_{y,0}$. According to Equations 2 and 3, the corresponding light field contains a single plane (perpendicular to $r_x$, $r_y$ and parallel to $k_x$, $k_y$) given by $r_{x,0} = P_x$ and $r_{y,0} = P_y$. According to the above analysis, axially displacing the point source only changes the orientation of this plane. Hence, $r_x = P_x$ and $r_y = P_y \; \forall P_z$. This is a simple statement of the fact that propagation alone does not change the direction of wave vectors constituting a wavefront - the ray perpendicular to the $\vec{k} = 0$ wavefront will intersect the first plane at $\vec{r} = r_{x,0}, r_{y,0}$ independently of the value of $P_z$.

Based on this analysis it is possible to considerably simplify Equation 1. Firstly, due to the symmetrical nature of the limiting apertures in most optical systems and further assuming this aperture is centred on the point $(k_{x,0}, k_{y,0}) = (0, 0)$ it can be seen that $c = d$:

$$ar_x + br_y + ck_x + ck_y + C = 0 \tag{4}$$

By setting the derivatives of Equation 4 with respect to $k_x$ and $k_y$ equal, it follows that $a = b$. Furthermore, translating the origin of the coordinate system to the intersect of the plane with the $r_x$ and $r_y$ axes, Equation 4 may be written as:

$$a\left(r_x - r_{x,0}\right) + a\left(r_y - r_{y,0}\right) + c\left(k_x + k_y\right) = 0 \tag{5}$$

Or, in vector notation:

$$a\left(\vec{r} - \vec{r_0}\right) + c\vec{k} = 0 \tag{6}$$

Defining disparity as $\vec{r_d} = \vec{r} - \vec{r_0}$ and $\alpha\left(P_z\right)$ as $\alpha\left(P_z\right) = -c/a$, it follows that:

$$\vec{r_d} = \alpha\left(P_z\right)\vec{k} \tag{7}$$

which, in Cartesian coordinates can be seen to have the form of a right circular cone with opening angle determined by the depth of the point in z:

$$x_d^2 + y_d^2 = \alpha^2\left(P_z\right)k^2 \tag{8}$$

#### B. Optical Imaging Model

The optical model used to perform simulations was designed to closely replicate the experimental system. The simulations were based on some key assumptions: monochromatic, isotropic fluorescent emission from point sources, no scattering and scalar diffraction. The pupil phase function due to a point source located at $(x, y, z)$ was defined as:

$$\begin{aligned}\mathcal{P}\left(k_x, k_y\right) = &P_{WF}\left(k_x, k_y\right)\exp\left[i2\pi\left(k_x\Delta x + k_y\Delta y\right)\right] \\ &\exp\left[i2\pi\left(k_x, k_y\right)\Delta z\right]\end{aligned} \tag{9}$$

where $(0, 0, 0)$ corresponds to a point source located on the focal plane, positioned at the optical axis. The point spread function of the widefield microscope is calculated as the two dimensional Fourier transform of this pupil function. In a light field microscope, this scalar wavefront is modified by a microlens array with pitch $(d_x, d_y)$ and focal length $f_{\mathrm{ML}}$. Applying the thin lens approximation, the phase modification due to a single microlens array:

$$T_{\mathrm{ML}}\left(x, y\right) = P_{\mathrm{ML}}\left(x, y\right)\exp\left[\frac{ik}{2_{\mathrm{ML}}}\left(x^2 + y^2\right)\right] \tag{10}$$

The full microlens array was generated by convolving this transmittance function with a two dimensional Dirac comb. The phase profile of the microlens array was multiplied with the widefield point spread function and finally propagated to the detector. Under the Fresnel approximation, the free space transfer function $\mathcal{H}\left(k_x, k_y\right)$ is:

$$H\left(k_x, k_y\right) = \exp\left[\frac{i}{4\pi}\lambda f_{\mathrm{ML}}\left(k_x^2 + k_y^2\right)\right] \tag{11}$$

Combining each of these equations, and noting that conventional sensors record irradiance, it follows that the light field point spread function may be computed as:

$$\mathrm{PSF_I} = \left|\mathcal{F}^{-1}\left[\mathcal{F}\{\mathrm{PSF_{A,\ WF}} \times \mathrm{T_{MLA}}\} \times H\left(k_x, k_y\right)\right]\right|^2 \tag{12}$$

The resulting images were scaled so that the integral matched the expected number of signal photons. Poisson noise was accounted for by replacing each pixel with a random number drawn from a Poisson distribution. Figure 9 shows a comparison between the experimental and simulated data.

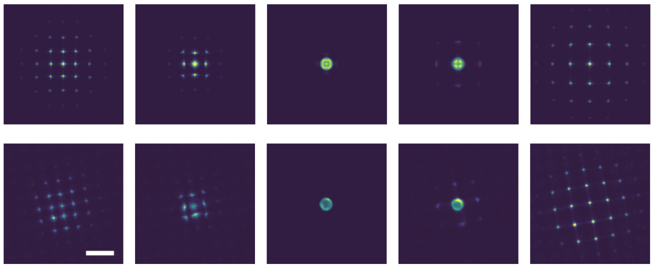

FIG. 9. Comparison between simulated point spread function (top row) and experimentally acquired data (bottom row). All simulations were performed using parameters chosen to match the experimental setup: pixel size 0.5 $\mu$m, ultimately downsampled to 6.5 $\mu$m pixels, microlens focal length 2.1 mm, pitch 100 $\mu$m, wavelength 525, $n = 1.4$, $M = 66.67$ and $NA = 1.3$. Scale bar indicates 2 $\mu$m.

#### III. SAMPLE PREPARATION

##### A. T Cell Labelling Protocol

Around $10^6$ T cells were labelled with 200 nM of the labelled TCR-(CAGE552) fabs (anti CD3, UCHT-1) on ice for 25 minutes. Cells were then washed three times in filtered PBS (involving centrifugation). Labelled T cells were fixed in 4% paraformaldehyde (Sigma) and 0.2% glutaraldehyde (Sigma) for 60 minutes at room temperature. The fixed cells were washed three times in filtered PBS and suspended in filtered PBS. For imaging, cells were stuck on a glass bottom dish and imaged using 561 nm excitation, with regular bursts of 405 nm activation laser.

##### B. Spermatocytes Labelling Protocol

Drosophila testes were dissected in ice-cold PBS, treated with collagenase (5 mg/mL; Sigma) for 5 min and cells were released by gentle pipetting. For fixing, cells were treated with PBS/4% formaldehyde and incubated for 20 min. Cells were then washed with PBS/0.01% Triton X-100 and filtered using 50 $\mu$m filter. The cells were resuspended in 10 $\mu$L PBS/0.01% then adhered to a glass bottom dish by slowly releasing them at the bottom of the dish, containing PBS, using a pipette.

The cells were blocked for 2 hours in PBS/0.5% WBR/0.5% Triton X-100 (WBR is Roche Western Blot Blocking Reagent) then incubated in primary antibody, MabE71 Mouse anti-Histone (1:1000 in PBS/0.5% WBR; Millipore). Cells were washed with PBS0.1% Tween-20). Secondary antibody solution, anti-mouse IgG conjugated with Alexa-647 (1:1000 in PBS/0.5% WBR; Invitrogen) was then added to the cells and incubated for 1.5 hours. The cells were washed with PBS /0.1% Tween-20 before fixation with PBS/4% formaldehyde for 20 min. Lastly cells were washed with PBS and stored at 4C. Imaging was carried out in Gloxy buffer, details of which can be found in [2].

[1] T. Peng, K. Thorn, T. Schroeder, L. Wang, F. J. Theis, C. Marr, and N. Navab, Nature Communications **8**, 14836 (2017).

[2] G. T. Dempsey, J. C. Vaughan, K. H. Chen, M. Bates, and X. Zhuang, Nature Methods **8**, 1027 (2011).
